# A novel approach to incorporate compound dissimilarity to plant chemodiversity measures using UV-Vis spectra

**DOI:** 10.64898/2026.08.24.746705

**Authors:** Kruthika Sen Aragam, Anke Steppuhn

## Abstract

1. Since plants interact with their environment through complex combinations of phytochemicals rather than metabolites in isolation, quantifying plant chemodiversity is of increasing interest. Although functionally relevant, structural disparity is mostly neglected in measures of chemodiversity because its integration relies on known compound identity, limiting broad application.
2. We established an approach deriving compound dissimilarity from UV-Vis spectra in HPLC-DAD datasets, evaluated how it relates to structure- or biosynthesis-based approaches, and examined whether it provides meaningful contributions to chemodiversity measures. For this, we applied it in an experiment full-factorially testing the effects of drought and herbivory on *Solanum dulcamara* leaf chemodiversity.
3. UV-Vis spectral dissimilarity aligned well with *fMCS*-based structural dissimilarity and reflected structural relationships within a set of standards. Incorporating it into chemodiversity analysis improved separation of the effects of drought and herbivory. Especially in the combination of both stresses, their distinct effects on different leaf metabolites were only fully reflected when accounting for compound disparity.
4. Hence, UV-Vis spectral dissimilarity captures chemically meaningful compound relatedness without requiring compound identity. Using it as a proxy for compound disparity adds a biologically relevant dimension to measures of chemodiversity with broader implications when assessing functional consequences of phytochemical diversity.

## Introduction

The extraordinary chemical complexity of plants, referred to as plant chemodiversity, is largely driven by secondary metabolites (also called specialised metabolites). Unlike primary metabolites, they usually are not required for basic metabolic processes but are often involved in maintaining plant development and survival in diverse interactions with their biotic and abiotic environment. Across the plant kingdom, at least one hundred thousand chemically unique compounds have been described so far (Kessler & Kalske, 2018; Forrister *et al*., 2023). An individual plant species can produce several thousand distinct secondary metabolites, with estimates in the range of five-to fifteen-thousand structures per species (Li & Gaquerel, 2021). Metabolic diversity differs not only between plant species but also within species, for example between genotypes and even within a plant individual across spatial (e.g. in different tissues) and temporal scales (e.g. over plant development or due to phenotypic plasticity).

Plant chemodiversity is assumed to be driven by the complexity of interactions that plants face in their environment (Iason *et al*., 2011; Dyer, 2018; Wetzel & Whitehead, 2020). In addition, various interacting effects between different metabolites such as synergisms and antagonisms entail that ecological functionality is determined by the collective properties of compound mixtures, which cannot be inferred from the effects of single compounds alone (Berenbaum & Neal, 1985; Steppuhn & Baldwin, 2007; Richards *et al*., 2015; Liu *et al*., 2017; Whitehead *et al*., 2021). Accordingly, recent studies indicate that chemical diversity can affect abundance and community structure of interacting species such as herbivores or pollinators. In addition to correlative evidence from studies that assessed metabolite diversity and herbivore communities in native plant populations (Richards *et al*., 2015; Volf *et al*., 2018; Salazar *et al*., 2018), there is experimental evidence from studies that manipulated chemodiversity at the level of plant communities using different chemotypes of a plant species (Bustos-Segura *et al*., 2017; Sasidharan *et al*., 2024). Consequently, there is growing interest in quantifying plant chemodiversity in standardised and comparable ways to enable testing how differences in the composition of metabolite profiles influence ecological interactions, plant resistance and fitness.

Different indices have been used to quantify phytochemical diversity at different levels of biological organisation such as cells, tissues, individuals, populations or communities and enable to examine how chemodiversity is constructed and is ecological relevant across these levels. In this regard, alpha, beta and gamma diversity are used to describe hierarchical diversity partitioning (Marion *et al*., 2015; Wetzel & Whitehead, 2020). While phytochemical diversity at a single sampling unit is defined as alpha-diversity, gamma-diversity is determined from pooled phytochemical profiles of sampling units at a higher organisational level and beta-diversity quantifies the variation between sampling units. As all approaches that investigate the role of chemodiversity in ecological and evolutionary processes depend on a meaningful quantification of diversity, they rely on metrics that integrate all relevant dimensions of chemical diversity.

The phytochemical composition of an individual plant forms a complex phenotype, which can be characterised along multiple dimensions such as the number of compounds (richness), the balance in their relative abundances (evenness) and the structural or biosynthetic dissimilarity of the compounds (disparity) (Marion *et al*., 2015; Junker, 2018). Most empirical studies quantifying chemodiversity apply classic diversity indices that combine richness and evenness such as Shannon, Simpson or Hill numbers (Bravo-Monzón *et al*., 2014; Li, 2018; Glassmire *et al*., 2019; Ziaja & Müller, 2023). These indices are straightforward to calculate from chromatographic data, as they do not depend on compound identity. Yet, they do not incorporate structural diversity of the analysed metabolites. However, as the molecular structure of a metabolite determines its biological activity, structural disparity among metabolites in a mixture can provide ecologically relevant information. In *Piper amalago*, for example, higher structural diversity significantly reduced herbivore feeding damage, whereas compositional diversity primarily explained variation in herbivore abundance and diversity, indicating that different dimensions of chemodiversity capture distinct ecological processes (Cosmo *et al*., 2021). In addition, structural disparity is indicative of the biosynthetic breadth underlying chemodiversity since phytochemicals that originate from shared biosynthetic pathways often share similar core structures. Both, the bioactivity of compounds and the complexity of the biosynthetic machinery producing them are of high ecological and evolutionary relevance, which raises the demand to include this previously neglected dimension, structural disparity, in frameworks to quantify chemodiversity (Petrén *et al*., 2024).

To date, only a small number of studies have explicitly quantified structural disparity in phytochemical mixtures. Initial attempts to account for structural disparity in chemodiversity measures were based on selected chemical characteristics, primarily chemical class or molecular weight. For example, Volf *et al*., (2018) calculated alkaloid and polyphenol diversity of *Ficus* species based on the contents of major structural groups rather than individual compounds, whereas Bakhtiari *et al*., (2021) used chemical properties as ‘functional traits’ to calculate multidimensional functional diversity indices according to Laliberté *et. al,* (2014) for glucosinolate profiles of Cardamine along elevational gradients. Approaches to incorporate structural disparity in a more standardised manner either rely on compound identification, so that structural similarity can be quantified based on molecular scaffolds or biosynthetic pathways, or on spectral similarity scoring of tandem-MS spectra (Richards *et al*., 2015; Sedio, 2017). The latter approach has been applied in metabolomic analyses of tropical Inga species which provided signatures of divergent adaptation, while a phylogenetic signal was lacking (Forrister *et al*., 2023). To test how structural diversity affects consumer responses, Whitehead *et al*., (2021) used synthetic mixtures of phenolic compounds to be able to apply scaffold- and biosynthetically informed metrics. Yet, investigating the role of structural disparity in naturally occurring metabolites relies on compound identification through NMR or MS/MS fragmentation spectra from which structural properties can be inferred (Cao *et al*., 2008; Cereto-Massagué *et al*., 2015; Wang *et al*., 2016; Ernst *et al*., 2019). This requires advanced and expensive analytical instrumentation. Acquisition and annotation of MS/MS or NMR spectra for all constituents in a mixture is labour intensive and often applied for selected metabolite groups (Li & Gaquerel, 2021). These limitations constrain the quantification of compound disparity and restrict the applicability of indices that include structural divergence in studies on the ecological and evolutionary roles of phytochemical diversity.

The approaches to quantify structural disparity are not applicable to other phytochemical profiling methods frequently used in ecological studies such as high-performance liquid chromatography coupled with diode-array detection (HPLC-DAD). It remains one of the most widely used methods as it is highly reproducible, affordable and relatively simple analytical technique for substance quantification. Although the detection does not provide structural information, the absorbance spectra of compounds reflect chromophore properties, which in combination with chromatographic separation are highly compound-specific. Consequently, HPLC-DAD has long been used as a highly reliable method for substance recognition, e.g. in toxicological analysis (Herzler *et al*., 2003). Yet, when HPLC-DAD profiles have been translated into chemodiversity measures, analysis remained restricted to include richness and evenness. For example, Almaraz-Ab *et al*., (2008) used retention times and UV-Vis absorbance spectra to differentiate phenolic compounds, but quantified variation among cactus taxa using Shannon diversity neglecting spectral (dis)similarity among compounds. Using a similar approach in a two-factorial experiment with milkweeds, Diethelm *et al*., (2024) detected an antagonistic effect between water limitation and herbivory on Shannon diversity of UV-Vis absorbing metabolites. The herbivory-associated increase in richness and the increased evenness under dry conditions disappeared if both stressors co-occurred, however compound disparity was not included in the analysis although very different metabolites that are differentially associated with these stresses were assessed (pregnane glycosides with herbivory, water stress with flavonoids). Consequently, these analyses did not capture whether the compounds contributing to richness and evenness are structurally similar or dissimilar.

Here, we explore the potential of using UV-Vis absorbance spectra to incorporate compound disparity into measures of phytochemical diversity from HPLC-DAD analysis. For this, we developed a workflow that derives compound dissimilarities from pairwise Pearson correlations between the UV-Vis absorbance spectra for each chromatographic peak. The resulting spectral similarity matrix is transformed into a dissimilarity matrix (**Fig. 1**) and subsequently used to evaluate different chemodiversity measures. We first assessed on a set of compounds of different chemical classes how structural dissimilarity based on UV-Vis spectral dissimilarity compares to existing methods and then evaluated whether the proposed approach to quantify structural disparity from HPLC-DAD analyses provides an ecologically meaningful component to measures of phytochemical diversity. To do so, we tested its ability to improve discrimination of phytochemical responses to two ecologically divergent stressors that are known to induce different phytochemical responses. These stressors, drought (abiotic) and herbivory (biotic), activate mainly distinct but interacting signalling pathways resulting in divergent and partially overlapping responses, and their combination can produce non-additive responses with consequences for herbivore performance (Nguyen *et al*., 2016, 2018). Based on this and previous studies (e.g. Diethelm *et al*., 2024), we expected an increased chemodiversity in response to both treatments and hypothesised that including compound disparity will separate treatment effects on chemodiversity more clearly than measures based only on richness and abundance. As the plant response to both treatments involve different metabolic responses, we further hypothesised that structural disparity especially contributes to differentiate a combined stress treatment from single stress treatments, as the metabolic changes induced by a single stressor may mask the effect of an additional stress when compound disparity is disregarded. Thus, we tested the approach to use UV-vis spectral dissimilarity as proxy for structural disparity in a full-factorial experiment with *Solanum dulcamara* plants with drought and herbivory as stresses and assessed leaf chemistry using HPLC-DAD.

**Figure 1:**
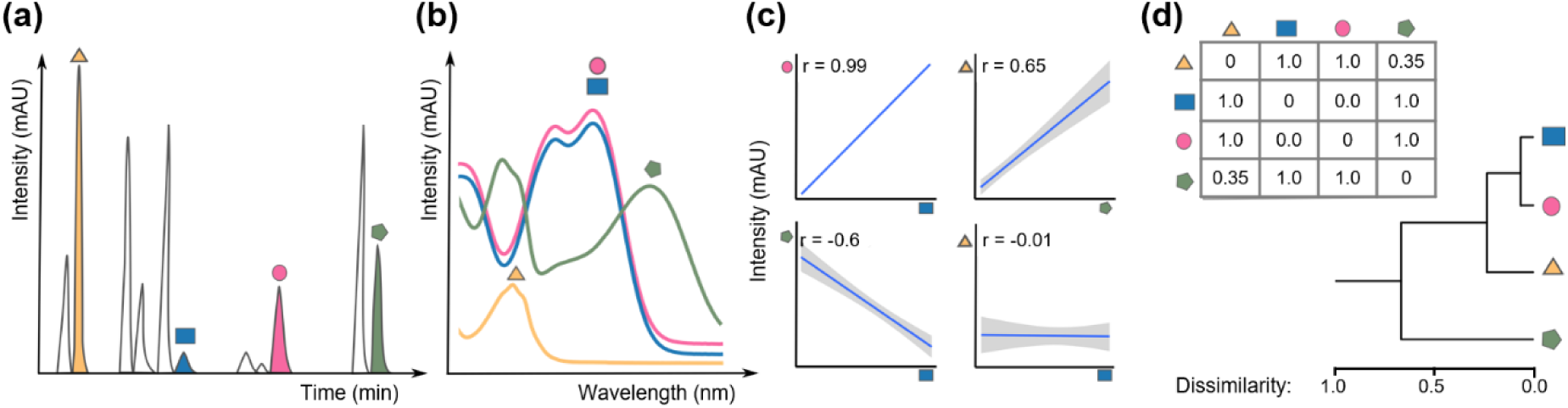
Workflow scheme for quantifying compound dissimilarity from UV-Vis spectra. Four example compounds are depicted by different shapes and colours in (a) chromatogram and their corresponding (b) UV-Vis absorption spectra, which are used for (c) pairwise Pearson correlations to determine spectral similarity. After conversion to dissimilarity, the (d) compound dissimilarity matrix is used for clustering of compounds into subclasses and to calculate diversity indices that incorporate structural disparity (i.e. FHD).

## Materials and Methods

### Plants & Insects

*Solanum dulcamara L.* plants of second-generation inbred lines derived from plants originally collected close to Ithaca, USA (GPS: 42°32’13.3“N 76°36’35.2”W) were used. Experimental plants were clones derived from 13 seedlings and each genotype was represented once in each treatment. Clones were propagated from stem cuttings that were allowed to root in 50mL centrifuge tubes for 1-2 weeks. Rooted cuttings were transplanted one node above and one below the soil into pots (1.1L, ⌀ 14cm) filled with fertilised (3g/L Osmocote Exact Standard 3-4M; Giulini GmbH, Germany) soil (*Substrate5* with clay + GreenFibre®; Klasmann-Deilmann GmbH, Germany). The soil surface was covered with a quartz sand layer (∼0.5cm). Plants were maintained pest-free using predatory mites (Amblyseius swirskii; Koppert GmbH, Germany) for biocontrol and were kept in a greenhouse (∼40% relative humidity; 22°C / 18°C during 16h light / 8h dark photoperiod).

*Spodoptera exigua* Hübner were reared as described earlier (Bandoly *et al*., 2015). In brief, colonies were maintained in a climate chamber (Polyklima GmbH, Freising, Germany) at 24°C with a 16:8h light:dark cycle. Larvae were fed *ad libitum* on a bean-flour based artificial diet in vented plastic rearing boxes and provided with paper tissue for pupation. Adults kept in flight cages (60x38x38cm) were supplied with 20% honey solution and paper tissue for oviposition. Egg clutches were collected for continued colony maintenance.

### Experimental Treatments and sampling

Three-week-old plants were randomly assigned to one of two watering regimes: well-watered (control) or drought. Soil moisture was monitored using an ML2x ThetaProbe connected to an HH2 Moisture Meter (Delta-T Devices). Control plants were watered every other day to maintain 35% volumetric water content (vmc) in soil. Plants assigned to the drought treatment were not watered until soil moisture declined to 5%vmc (after six days), and were maintained at this level for the experimental period by suppling minimal water amounts according to daily soil moisture measures (**Fig. S1**). Seven days after drought was initiated, all plants were sampled for leaf tips of two young source leaves (4 and 5 positions below the visually predicted sink-source transition leaf) and herbivory was initiated on the same leaves. One third-instar larva was placed per leaf and confined using fine-mesh bags. Larvae were allowed to feed for 72h, before mesh bags and larvae were removed and the leaves were resampled. At each harvest, the samples of the two leaves were pooled and immediately flash-frozen in liquid nitrogen and stored at -80°C until further processing. Thus, the effect of herbivory was assessed from the same leaves before and after herbivory, in a paired design, while drought was applied on one of two clones for each of the 13 genotypes.

### Leaf extraction and HPLC-Analysis

An aliquot of the leaf material of each homogenised pool was used to determine dry matter content by measuring its weight before and after drying at 80°C for overnight in 1.5mL tubes. For extraction, approximately 100 (± 20) mg of each pool were transferred in 2mL screw-cap tubes pre-filled with ceramic beads (2.8-3.2mm; Zirconox, Mühlmeier GmbH & Co. KG, Germany) and the exact fresh weight was documented. After homogenisation in 1 mL of 40% aqueous methanol containing 0.5% acetic acid using a Bead-Mill24 (Fisherbrand™, FisherScientific GmbH, Germany) for 2 x 20s at 4ms⁻¹ and 15min of vortexing, extracts were centrifuged at 15,000g for 15min at 22°C. The supernatant was transferred to 1.5mL tubes and upon a second centrifugation (15,000 x g for 10min), 200µL of the clarified extract was transferred into HPLC glass vials with inserts and stored at 4°C until HPLC-DAD analysis.

HPLC-DAD (Shimadzu SPD-M20A, Tokyo, JP) instrument with a binary pump, a vacuum degasser, an autosampler, a column compartment, and a diode array detector was used. Reverse phase, C18-column (Inertsil ODS 3μm, 3.0 x 250mm) attached to a C-18 precoulmn was used for compound separation at 28°C using 0.25% H3PO4 in water as eluent A and acetonitrile as eluent B. Compounds were analysed from 10µL of the extracts at 0.5ml/min flow in gradient mode (eluent B set at 0min: 6%, 2min: 6%, 6min: 9%, 20min: 18%, 41min: 28%, 44min: 33.3%). Between runs the system was rinsed and subsequently equilibrated (5min at 100% and 6min at 6% B). The retention times and compound-spectra were used to characterise 35 different compounds, among them chlorogenic acid, caffeoylputrescine and rutin that were identified by comparing compound spectra with the injections of authentic standard compounds (see **Tab. S11** for CAS no. and suppliers all used standards). Compound abundance was quantified from the peak areas at different wavelengths depending on the compound-specific absorbance maxima (**Tab. S1**) and expressed relative to the extracted leaf dry weight that was calculated from fresh weight based on the dry matter content of an aliquot of the same leaf sample.

### Quantifying compound dissimilarity from UV-Vis spectral data for integration in diversity measures

Spectral data of each of the reference compounds and all peaks analysed in the experimental samples (from chromatograms with high peak abundance) were exported from the HPLC-instrument software (LabSolutions, Shimadzu, Tokyo, JP). The three-dimensional chromatograms (absorbance intensity over retention time and wavelengths) were imported into R using the *read_chroms()* function of the R package CHROMCONVERTER (Bass 2024). The spectral data were reduced to a wavelength range of 240-380nm retaining only compound-specific absorbance characteristics of all analysed peaks. None of the detected compounds absorbed wavelengths above 380nm and absorbance below 240nm was dominated by nonspecific background absorbance. For each chromatographic peak included in the analysis, a representative chromatogram in which the peak occurred at high abundance and its corresponding retention time were selected manually (**Tab. S12**). Using the R-function *cor()*, Pearson correlations are then calculated between the UV-Vis absorbance spectra or all possible pairs between chromatographic peaks to generate a spectral similarity matrix. Similarities within the chromatophores of compounds are reflected in positive correlations between their absorbance spectra, yet negative correlations between absorbance spectra (e.g. when shifts of absorbance maxima result in opposing slopes of the spectra) are not indicative of greater structural disparity. Therefore, only positive Pearson’s coefficients are used for the correlation matrix (c), and negative values are set to 0. The similarity matrix is then converted to a dissimilarity matrix (d) by the formula: d_ij_ = 1-c_ij_ (where *i* and *j* are two individual compounds). This compound dissimilarity matrix is then used for the evaluation of chemodiversity employing the Rpackage CHEMODIV (Petrén *et al*., 2023), which enables quantification of multiple dimensions of chemodiversity, including compound disparity through functional Hill diversity (FHD) index. Both, Hill diversity (HD) and FHD were computed for diversity orders q0-q5 using the functions *calcDiv()* and *calcDivProf()*.

For six authentic standard compounds, pairwise compound dissimilarity matrices were additionally quantified based on three existing methods using the Rpackage CHEMODIV. All are based on calculating Jaccard dissimilarities between pairs of compounds from SMILES strings and InChIKeys (obtained from PubChem database) as input. The first method based on the deep-learning tool NPClassifier (Kim *et al*., 2021; NPC) is largely a measure of the biosynthetic dissimilarity of two compounds. Two other methods are based on structural dissimilarity, either using PubChem molecular fingerprints (PCmf) (Cereto-Massagué *et al*., 2015) or largest substructure that occurs in two metabolites using the graph-based flexible Maximum Common Substructure (fMCS) method (Cao *et al*., 2008; Wang *et al*., 2013).

### Statistical analysis

All analyses were performed in R version 4.4.0 (2024) and all data visualisation and statistical summaries were produced using the ggPLOT2 and RMISC packages. Based on the dissimilarity matrices, compounds were hierarchically clustered with average linkage using the DENDEXTEND R package. To evaluate the method of deriving compound dissimilarity from UV-Vis absorbance spectra in relation to existing methods, pairwise correlations among compound dissimilarity matrices were evaluated using Mantel tests, implemented with the function *mantel* (VEGAN package).

The experimental data set was analysed for treatment effects of drought and herbivory on single compounds as well as on HD- and FHD-indices using linear mixed-effects models (LMMs) implemented with the function *lmer* (LME4 package). Untransformed values were analysed as QQ plots and Shapiro-Wilk tests indicated normality and Levene’s test (CAR package) homogeneity of variances. Models included drought, herbivory, and their interaction as fixed factors and genotype as a random factor and accounted for the repeated measure. If an interaction was indicated at p < 0.1, contrasts for the effects of drought and herbivory in absence and presence of the other was tested separately using estimated marginal means. All p-values for the treatment effects on single compounds were corrected for false discovery rates (FDR) according to Benjamini-Hochberg.

Compound contributions to treatment dissimilarities were evaluated using SIMPER analysis (function simper, VEGAN package), based on Bray-Curtis dissimilarities derived from compound relative abundances. Average contributions and associated p-values were used to identify compounds primarily responsible for variation between treatments.

## Results

### Method comparison of compound dissimilarity metrics using authentic standards

UV-Vis spectral dissimilarity was compared to three different existing methods for quantifying pairwise dissimilarity of authentic standard compounds covering several chemical subclasses: two phenol amides, a phenolic acid, a flavonoid, and two alkaloids (**Fig. 2a**). All approaches reflected the close structural relationship between the two phenol amides, yet structural relatedness of the remaining compounds was rated rather differently as visualised by hierarchical clustering applied to each of the four dissimilarity matrices (**Fig. 2b**, **Tab. S4**). While equally dissimilar using *NPClassifier*, these compounds were more clustered with the other approaches. Only using UV-Vis spectra and *flexible maximum common substructure* (*fMCS*), the close structural relationship between the phenolic acid and phenol amides was reflected, while the latter were grouped with the structurally less similar alkaloid colchicine based on *PubChem molecular fingerprints* (*PCF*). *PCF*-derived compound dissimilarities grouped the phenolic acid more closely to the flavonoid rutin, reflecting rather a biosynthetic relationship than structural similarity. The most significant correlation in the Mantel test was found between UV-Vis spectral and *fMCS-*based dissimilarity (p = 0.007). The latter also correlated with the *PCF* (p = 0.013) and *NPClassifier* (p = 0.044), but UV-Vis spectral dissimilarity did not correlate significantly with these matrices (**Tab. S5**).

**Figure 2:**
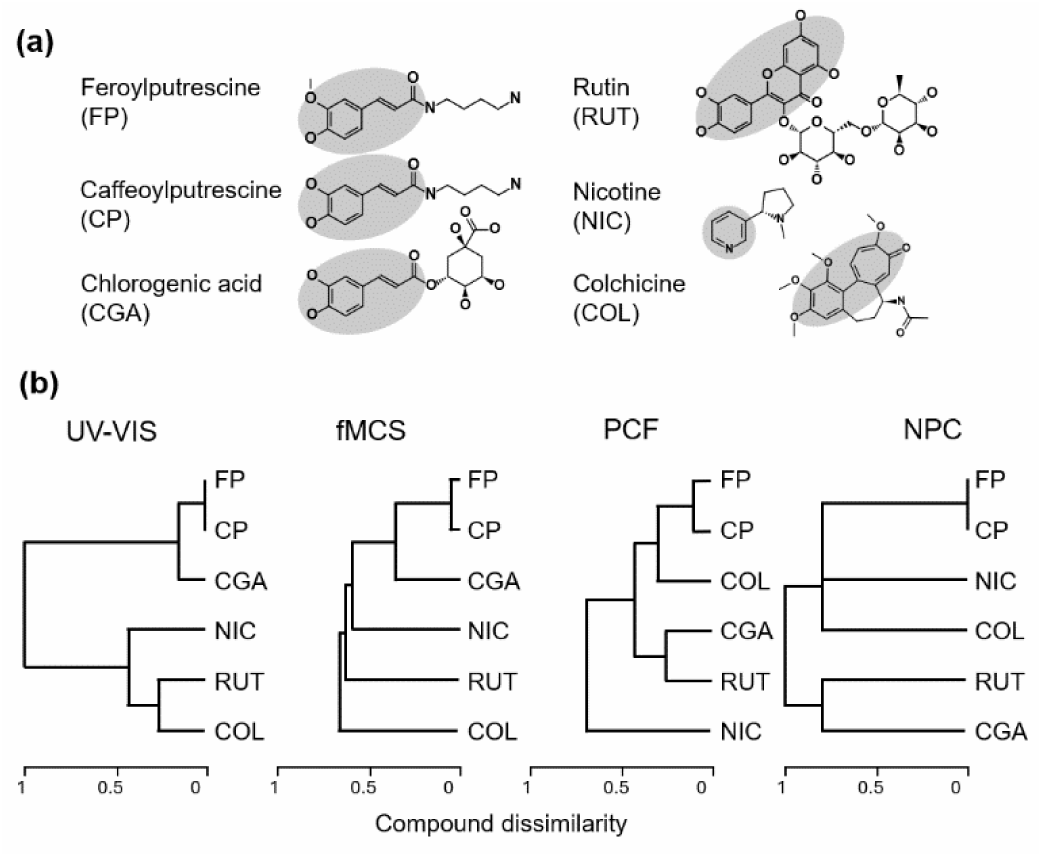
Structural dissimilarity of authentic standard compounds. (a) Chemical structures with highlighted chromatophores (drawn with PubChem Sketcher) and (b) hierarchical clustering using average linkage based on dissimilarity matrices derived from UV-Vis spectra in comparison to flexible maximum common substructure (fMCS), PubChem fingerprints and NPClassifier. Branch lengths in the dendrograms represent relative dissimilarity among compounds.

### Absorbance based dissimilarity of compound in *S. dulcamara* leaves

In the leaf extracts of *S. dulcamara* plants in the full-factorial experiment with drought and herbivory treatments, we detected 35 compounds with quantifiable peaks in no less than a third of the samples in at least one of the four treatment groups. Four of them were determined as the two phenol amides caffeoyl- and ferroylputrescine (CP and FP), chlorogenic acid (CGA), and rutin (RUT) by matching retention times and absorbance spectra with the authentic standards. According to UV-Vis spectral dissimilarity, two major clusters with less than 20% spectral dissimilarity contained 75% of the compounds, among them are FP, CP, and CGA (**Fig. 3a**). The remaining quarter consisted of one compound clustering with rutin, two compounds with high dissimilarity to all others and a cluster of five (grey in **Fig. 3a**), in which one that was slightly separated from the others showed greater similarity to the spectrum of nicotine, which could indicate presence of a pyridine ring. Thus, richness of compounds with chromatophores in *S. dulcamara* leaves is largely driven by phenolic compounds (three lower clusters in **Fig. 3a**), yet at another dimension of chemical diversity, the abundance, other compound groups were important contributors. Peak intensity varied strongly between the compounds (**Fig. 3b**) with the five most abundant peaks contributing around three quarters of the sum of all peak areas in a chromatogram. These included CGA and a likely isomer (RT 17.7) with an almost identical absorbance spectrum to CGA, one of the very distinctly absorbing compounds (RT 40.1) and one in the first cluster (RT 02.3). Only one of the five most abundant peaks belonged to the cluster containing the two phenol amides (CP, FP). Mean relative peak abundance of 20 peaks were below 1 % of the total summed peak area. Rutin, CP, and FP were among compounds with an intermediate relative abundance (1-3 %), which were also dominated by compounds in the cluster with CGA. Both, richness and abundance of compounds, were affected by the drought and herbivory treatments.

**Figure 3.**
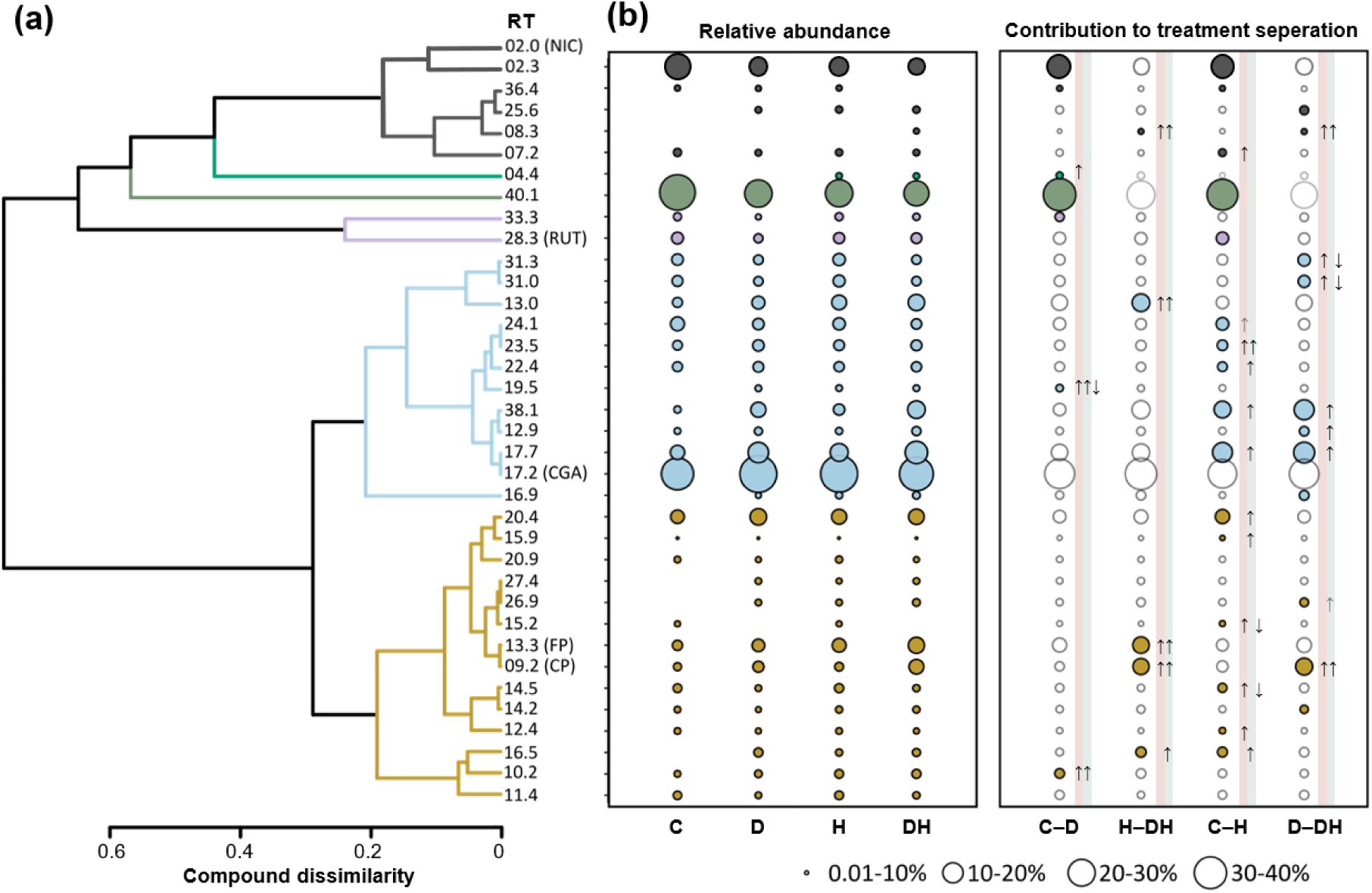
Dissimilarity of UV-Vis spectra of *Solanum dulcamara* leaf compounds and their relative abundance and contribution to discriminate drought and herbivory treatments. a) Dendrogram according to hierarchical clustering using average linkage based on pairwise UV-Vis spectral dissimilarity among compounds that are labelled by retention time (RT) in minutes. Clusters linked at dissimilarities below 0.25 are coloured differentially. Authentic standard injections revealed presence of caffeoyl- and ferroylputrescine (CP and FP), chlorogenic acid (CGA), rutin (RUT) but not of nicotine (NIC). b) Circle diameters in the left box represent mean compound abundance (relative to total peak area sum) in untreated to control (C) plants, and plants treated by drought (D), herbivory (H), or both (DH). Circles in the right box denote mean contribution of compounds to Bray-Curtis dissimilarity between treatment pairs according to SIMPER analysis. Circles are filled according to clusters in the dendrogram. Compound contribution is only filled if significant (p < 0.05) and then direction of regulation by drought and herbivory according to Fig. 4 is signified by arrows on red and green shading, respectively.

### Effects of drought and herbivory on leaf compound composition

We first analysed the effects of drought, herbivory, and their interaction on abundance of individual leaf compounds. Both treatments altered compound abundance with overlapping and interacting effects, which can be grouped into four distinct regulation patterns (**Fig. 4a,b**, **Tab. S8**). While twelve compounds were significantly increased by one of the treatments, either drought or herbivory (three of group I and all in IV in **Fig. 4**), fourteen compounds were affected by both treatments but in different ways. For nine of these compounds (group II), an antagonistic interaction between both treatments was indicated (for six significantly, for three by trend) because they were significantly increased in abundance by drought, while but this increase was supressed after herbivore exposure. Yet, only for two of them the reduction of drought-induced levels due to herbivory was significant (RT 19.5 and CGA). The other five compounds (group III) affected by both treatments were additively increased in their abundance in response to both treatments. For two compounds even a synergistic induction by the combination of drought and herbivory was indicated either by trend (RT 08.3: p = 0,057 after FDR correction) or significantly for the phenylpropanoid CP. With only one exception (RT 08.3), all compounds in groups II-IV were clustered with CGA or the phenol amides CP and FP and compounds with divergent absorption spectra were only responding to drought.

To assess the contribution of the individual compounds to treatment effects on the overall compound composition, we analysed similarity percentage (SIMPER) partitioning. Across all contrasts, relatively few compounds accounted for most of the between-treatment cumulative dissimilarity (**Fig. 3b**; **Tab. S7**). Yet, significance of the contribution of compounds was not associated with high relative contribution to treatment dissimilarity and thus abundance of compounds. Although only five to seven compounds accounted for ∼75% of cumulative dissimilarity between treatments, only half of these were significantly contributing for most contrasts. Only for the effect of herbivory in the absence of drought (C-H contrast), contributions of five of the six top-ranked compounds explaining 75% of the treatment separation were significant (and permutation p-value of that of second-ranked CGA was 0.093). Interestingly, a far lower number of compounds contributed significantly to the separation of drought treatments (7 and 5 compounds to C-D and H-DH contrasts) than to the separation of herbivory treatments (16 and 11 compounds to C-H and D-DH contrasts), although drought and herbivory affected abundance of a similar number of compounds (**Fig. 4** group I-III and group II-IV, respectively). As a consequence, cumulative between-treatment dissimilarity of significantly contributing compounds strongly differed between contrasts and was much higher for herbivory (C-H: 66%, D-DH: 33%) than for drought contrasts (C-D: 43%, H-DH: 18%).

Together these analyses show that the compounds significantly contributing to treatment differences in overall compound composition are not identical with the set of compounds affected by the treatments in their abundance. Especially in contrasts to control plants, treatment dissimilarities in composition were also driven by compounds that were themselves not affected, while many compounds significantly affected in abundance did not contribute to overall compound composition. Apart from that, the compounds only regulated by herbivory in **Fig. 4** (group IV) were the most dominant group of regulated compounds among the significantly contributing compounds to herbivory contrasts in compositional changes.

**Figure 4.**
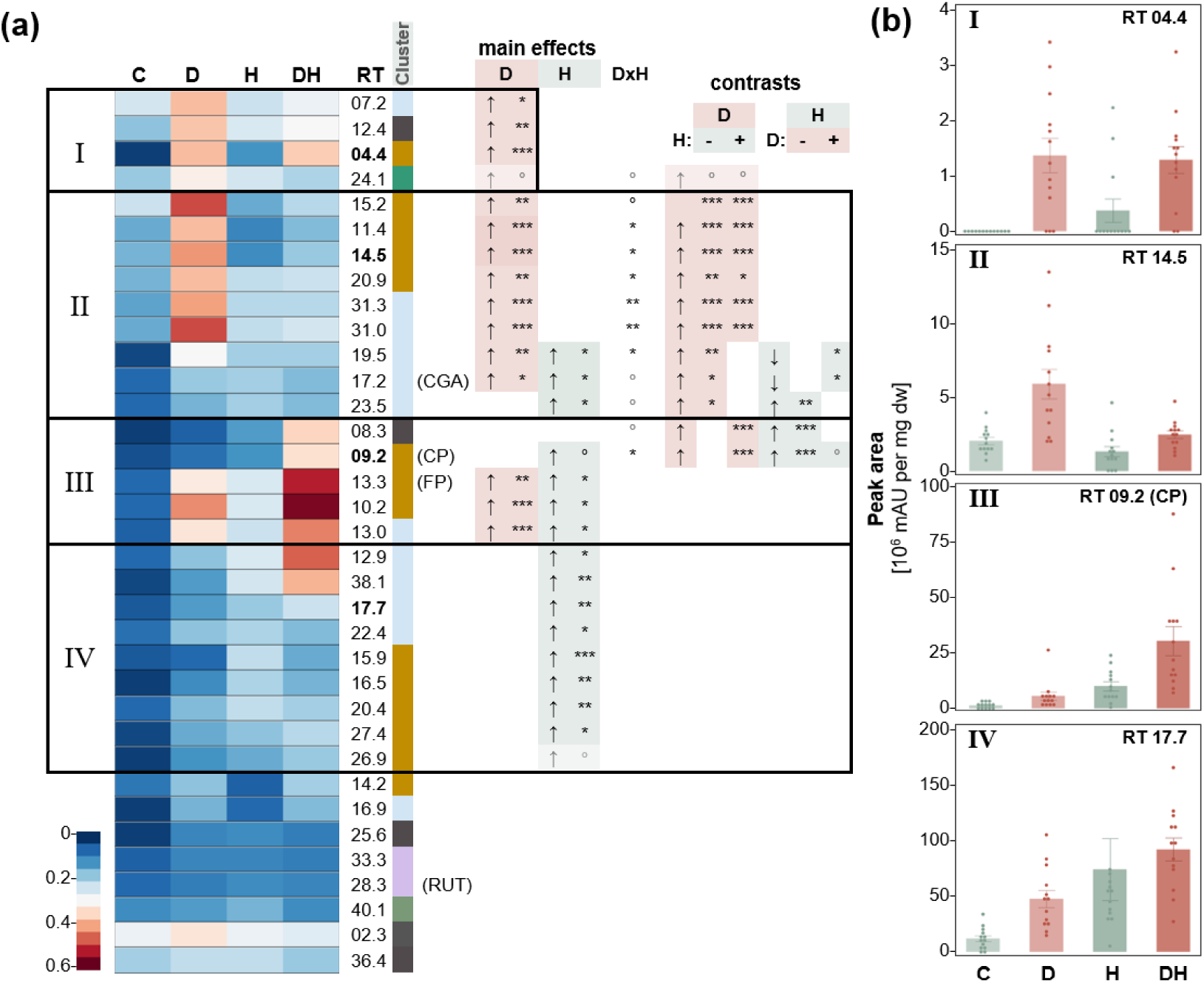
Treatment effects on individual compound abundance in *Solanum dulcamara* leaves of untreated to control (C) plants, and plants exposed to drought (D), herbivory (H), or both (DH). (a) Heat map for mean abundance (min-max normalised peak area) of compounds (labelled as in Fig. 3 and sorted for treatment effects). For each compound, the effects of D, H and their interaction (DxH) were tested with linear models (REML) and significant effects are indicated at the right. If an interaction was indicated (p < 0.1), pairwise contrasts (based on estimated marginal means) were used to test the effects of D and H in the absence and presence of the other factor. All p-values were FDR corrected according to Benjamini-Hochberg. (b) Peak areas (single data points and bars for mean +/-SE) of example compounds within groups distinctly affected by treatments (groups I - IV).

### Effects of drought and herbivory on chemodiversity incorporating structural dissimilarity or not

Compound richness of *S. dulcamara* leaves, corresponding to Hill diversity at order 0 (HD_q0_), increased significantly due to drought and herbivory (**Fig. S2**, **Tab S9**). The increase in number of compounds by the combination of both treatments was less strong than when applied as single treatments resulting in a significant interaction term (DxH) for HD_q0_. This interaction is lost when including structural dissimilarity based on UV-Vis spectra, because the effect of both treatments was rather additive resulting in even higher FHD_q0_ in plants of the combined treatment compared to the already elevated FHD_q0_ of plants of the single treatments. Although, evenness at q1 was not significantly affected by the treatments, it tended to be increased by drought (**Fig. S2**). At higher diversity orders, drought and herbivory significantly increased chemodiversity quantified as HD (**Fig. 5a**; **Tab S9**). Including structural dissimilarity (in FHD), strongly increased treatment separation especially at higher diversity orders (**Fig. 5b**; **Tab S9**). Significance level of treatments on HD rapidly dropped with increasing diversity order but remained relatively stable for FHD over diversity orders. This was more pronounced for the effect of herbivory compared to for that of drought resulting in a significant interaction for FHD from q3 on.

**Figure 5:**
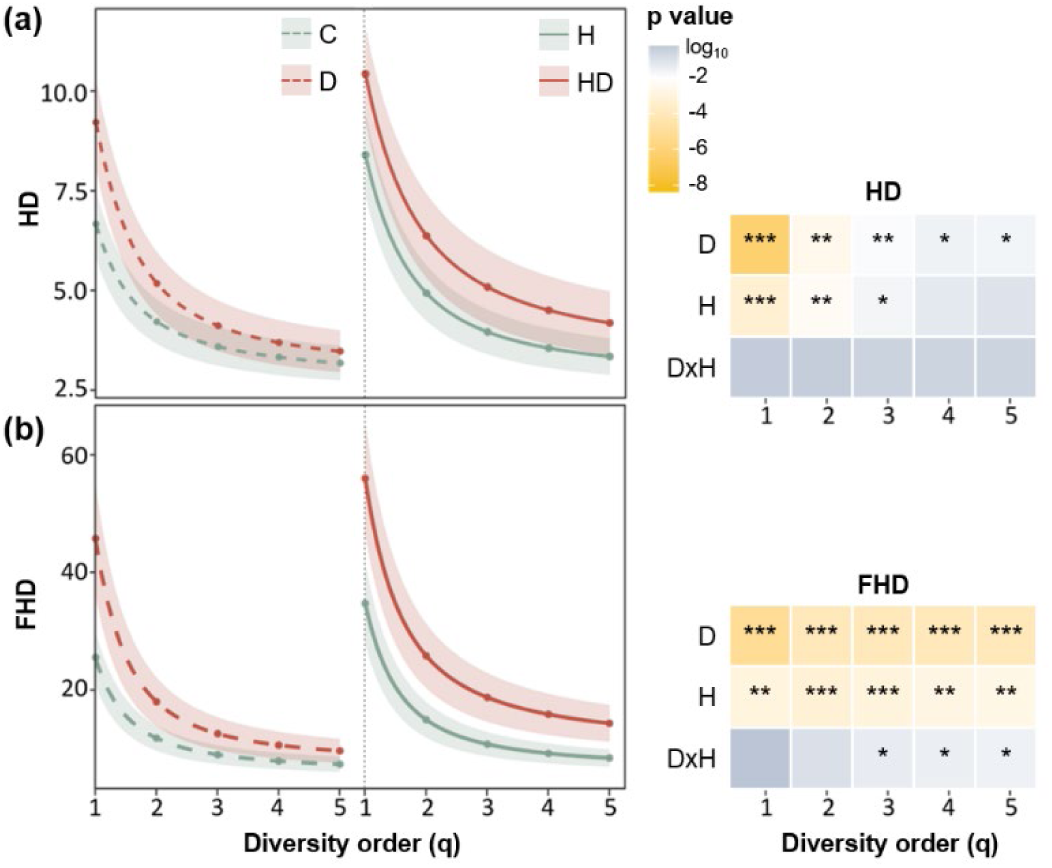
Chemical diversity profiles over diversity orders in *Solanum dulcamara* leaves of untreated to control (C) plants, and plants exposed to drought (D), herbivory (H), or both (DH). Lines depict means and shaded areas 95% confidence intervals of (a) Hill diversity (HD) and (b) functional Hill diversity (FHD). Right of the graphs treatment effects according to two-factorial linear mixed models for the factors D and H are depicted in table cell colours according to the p-values. Symbols (***/**/*) denote significant differences at p < 0.001/0.01/0.05 respectively.

## Discussion

Comparison of UV-Vis spectral dissimilarity between authentic standard compounds covering different chemical subclasses with three existing methods for quantifying pairwise dissimilarity revealed strongest correlation with *fMCS*-based dissimilarity. The lack of a such a correlation with dissimilarity derived from *NPClassifier* could be due to an overall low differentiation by the latter method for the structural dissimilarities of the analysed compounds. An exception was the very close structural similarity between the two phenol amides, which was consistently reflected in the dissimilarity matrices of all approaches (**Fig. 2b**). The low correlation between UV-Vis and *PCF* dissimilarity matrices likely results from two major differences. Firstly, the low *PCF*-based dissimilarity between colchicine and the structurally rather different phenol amides. Secondly, a low UV-Vis spectral dissimilarity between the two alkaloids and the flavonoid, which is likely driven by the fact that absorption is only determined by those parts of a chemical structure with conjugated π-electron systems (due to conjugated double-bonds such as in aromatic rings and attached functional groups that act as auxochromes), at which these compounds do show commonalities (**Fig. 2a**). The close structural relationship between CGA and the two phenol amides was only reflected in UV-Vis spectral and *fMCS* derived dissimilarities, likely contributing to the much better correlation between these two approaches. The *fMCS* method, which compares atom numbers in common substructures to the atom number of the whole molecule, has been suggested to have an increased performance to measure their structural dissimilarity (Wang *et al*., 2013). This method correlated with all other methods but most significantly with UV-Vis spectral dissimilarity indicating that compound dissimilarity derived from UV-Vis spectra can capture chemically meaningful aspects of compound relatedness.

Based on UV-Vis spectral dissimilarity, 70% of the light absorbing compounds in *S. dulcamara* leaves to compounds are closely related to phenolic acids and their derivatives such as phenol amides (**Fig. 3a**). Phenolic acids such as CGA were among the most abundant compounds. However, most of the compounds with related spectra were low and intermediately abundant. Although fewer in number, compounds with very divergent absorption spectra contributed on average to 50% of the total peak area in untreated control plants. However, this proportion almost halved in response to herbivory and drought treatments, because both treatments merely, in case of herbivory almost exclusively, induced compounds related to phenolic acids or phenolic amides. While three quarters of these compounds responded specifically to either drought or herbivory, only six compounds were induced by both. Among those were CGA and the phenolic amides FP and CP and for the latter a synergistic induction by both treatments was evident (**Fig. 4**). The especially strong inducibility of phenolic amides by both treatments is consistent with previous studies and their role in plant resistance against herbivores and drought in Solanaceous plants has been shown (Onkokesung *et al*., 2012; Cao *et al*., 2024). Apart from CP and one other compound (RT 08.3), all other compounds for which an interaction between drought and herbivory was indicated significantly or by trend (compounds in group II in **Fig. 4**), showed an antagonistic effect. For most of these compounds, drought induction was substantially lower in the combined treatment. Thus, consistent with previous literature (Suzuki *et al*., 2014; Nguyen *et al*., 2016, 2018; Mundim & Pringle, 2018), each of the two stresses induced largely divergent responses on UV-absorbing leaf metabolites in *S. dulcamara* and their combination alters compound composition differently than the single stresses.

The SIMPER analyses of the treatment effects on the overall compound composition revealed that more compounds contributed significantly to compositional changes in response to herbivory than in response to drought (**Fig. 3**), even though both treatments affected abundance of approximately equal numbers of compounds. However, the set of compounds significantly contributing to treatment separation in the overall composition overlapped by only about one third with the compounds significantly regulated by the respective treatment. The proportion of treatment-regulated compounds contributing to compositional changes was higher for herbivory than for drought (10 and 7 of 16 regulated by each stress). Abundant but non-regulated compounds contributed mainly to contrasts between treated and control plants. This can be explained by their lower relative abundance after treatment, as total peak area in control plants was only about half that of the treatment groups. Therefore, the induction of many low abundant peaks, even if not significant for individual compounds (for example if pathways produce sets of different metabolites, which can increase variation of the induction effect for a single metabolite) can be evident in compositional changes from the reduced relative abundance of unregulated but abundant metabolites. Consistent with that, in contrasts between single and combined treatments, which did not differ in overall peak area sum, compounds significantly contributing to compositional changes were dominated by regulated compounds.

Both, regulated and composition-affecting compounds, were widely spread across the dendrogram depicting spectral dissimilarity as an approximation of structural disparity (**Fig. 3**). To test whether this approach can provide meaningful and ecologically relevant contributions to measures of chemodiversity, we hypothesised that including spectral dissimilarity would improve treatment separation. Since, both treatments result in strong compositional changes that are also driven by increasing sets of regulated compounds relative to abundant ‘housekeeping’ metabolites and because the two divergent stressors induce distinct, overlapping and interacting effects, we especially expected a better separation of single versus the combined stress treatment on measures of chemodiversity. The treatment effects on single compounds and compound composition resulted in increased chemodiversity at different levels. While compound richness of (HD_q0_) was equally higher in drought and herbivory-treated plants compared to control plants, evenness tended to be only affected by drought (p = 0.093, **Fig. S2a**). These results in *S. dulcamara* are very similar to those of Diethelm *et al*. (2024) assessing the effects of herbivory and water limitation on milkweed chemodiversity. Similar to this study, we also observed an antagonistic interaction between drought and herbivory on richness (HD_q0_). However, in contrast to the loss of the herbivory-associated increase of milkweed metabolites under water limitation, the combined treatment in *S. dulcamara* affected compound richness less strongly but still increased it. Including spectral dissimilarity as a measure of disparity based on UV-Vis spectra, suggested that the apparent antagonistic response was associated with not considering compound disparity, as the effects of drought and herbivory were approximately additive for FHD_q0_. Moreover, FHD strongly improved treatment separation throughout diversity orders (**Fig. 5b**), which was particularly pronounced at higher diversity orders, progressively assigning greater weight to compound abundance. This was especially observed for the effect of herbivory on chemodiversity, as herbivory predominantly affected less abundant peaks. At higher diversity orders, including the dimension of compound disparity revealed a synergistic interaction between drought and herbivory, particularly in contrasts between the combined and single-stress treatments (**Fig. S2b**). Together, these results indicate that including structural disparity provides a more comprehensive assessment of chemodiversity responses, particularly where effects of combined stresses are not apparent from richness and evenness alone.

## Conclusions

Chemodiversity is increasingly used to describe complex phytochemical phenotypes, but structural disparity remains much less commonly considered than richness and evenness mainly because structural diversity quantification often relies on cost- and labour-intensive analytical approaches. Here, we show that UV-Vis spectral dissimilarity derived from HPLC-DAD analyses provides a practical approximation of structural disparity for compounds with UV-Vis absorbing chromophores and corresponds particularly well with *fMCS*-based estimates, which is regarded as one of the most powerful methods. We further demonstrate the ecological relevance of this approach in a drought and herbivory experiment with *S. dulcamara*, where including UV-Vis spectral dissimilarity improved separation of single and combined stress treatments. Thus, the approach provides a comparatively simple way to integrate structural disparity into chemodiversity measures when detailed structural identification of all compounds are not available.

## Acknowledgements

We thank the German research foundation DFG for funding under the project number 415496540 (STE2014/6-1) of FOR3000. We sincerely thank Lu Li for her valuable inputs during experimental design planning, implementing drought treatment and supporting with plant cultivation. We also thank the students of experimental plant ecology course (1901-240, 23/24) at Uni Hohenheim, for their assistance with sample extraction and herbivory treatment. We are grateful to Jens Schwachtje for his thoughtful comments on the manuscript.

## Author contributions

AS conceptualised the approach with input from KSA. KSA carried out the experimental work, including the sample extraction, developed the analytical workflow, performed chemical and statistical analysis. KSA and AS jointly wrote and edited the manuscript.

## Data availability

All data, supporting information and Rmarkdown files will be deposited on DataPlant for public access.

## Supporting information

**Fig. S1:**
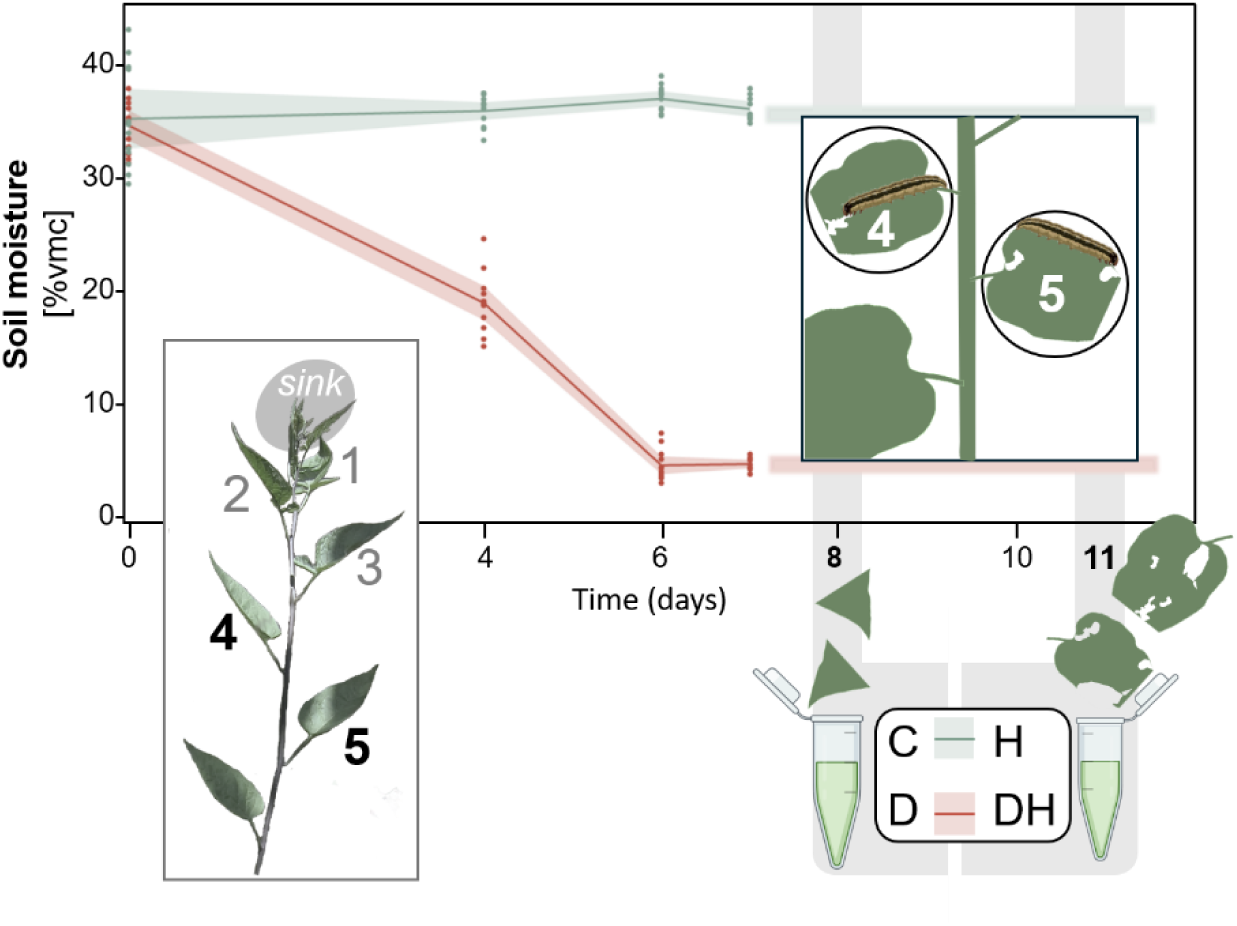
Soil moisture and experimental timeline for drought and herbivory treatments. Drought was applied by withholding water to over a week, in which relative volumetric moisture content (% vmc, mean ± SD) of soil was recorded (on day 0, 4, 6, 7) of well-watered and drought treated plants. Within six days, soil moisture of drought treated plants declined to about 5% vmc, which was maintained at this level throughout the experiment by low water supply according to soil moisture measurements, while control plants were supplied with excess water and maintained soil moisture around 35% vmc. On day 8, the tips of two leaves (4th and 5th youngest source leaves) were sampled and pooled to assess the effect of drought in the absence of herbivory (control: C and drought: D treatment group) and on both leaves one 3rd instar *Spodotera exigua* larvae was enclosed using mesh bags. On day 11 larvae were removed and the leaves were sampled without midribs to assess the effect of herbivory in the absence and presence of drought (herbivory: H and drought + herbivory: DH treatment group). An aliquot of each pooled sample was assessed for leaf water content by taking weights before and after drying and the remaining sample was weighted and extracted for HPLC analysis.

**Fig. S2:**
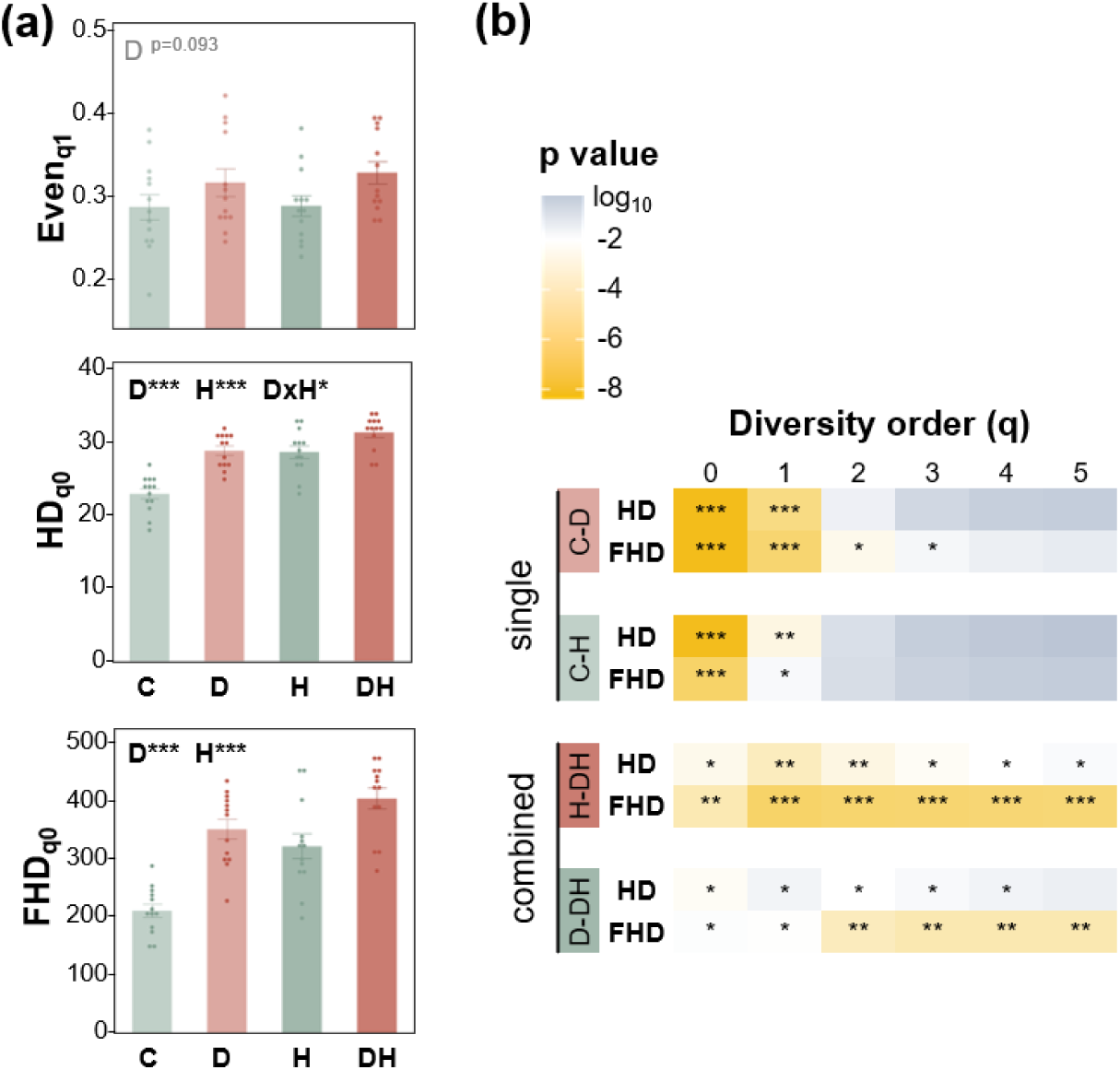
Chemical diversity in *Solanum dulcamara* leaves of untreated to control (C) plants, and plants exposed to drought (D), herbivory (H), or both (DH) and significance of pairwise contrasts throughout diversity orders. (a) Evenness (Even_q1_) and compound number corresponding to Hill diversity (HD) at diversity order 0 and Functional Hill diversity (FHD_q0_) incorporating compound disparity to richness based on UV-vis spectral dissimilarity. (b) Significance of pairwise contrasts throughout diversity orders.

## References

Almaraz-Ab N, Campos MG, Delgado-Al EA, Avila-Reye JA, Herrera-Co J, Gonzalez-V LS, Naranjo-Ji N, Frigerio C, Tomatas AF, Almeida AJ, et al. 2008. Pollen flavonoid/phenolic acid composition of four species of Cactaceae and its taxonomic significance. American Journal of Agricultural and Biological Sciences 3: 534–543.

Bakhtiari M, Glauser G, Defossez E, Rasmann S. 2021. Ecological convergence of secondary phytochemicals along elevational gradients. New Phytologist 229: 1755–1767.

Bandoly M, Hilker M, Steppuhn A. 2015. Oviposition by *Spodoptera exigua* on *Nicotiana attenuata* primes induced plant defence against larval herbivory. The Plant Journal 83: 661–672.

Berenbaum M, Neal JJ. 1985. Synergism between myristicin and xanthotoxin, a naturally cooccurring plant toxicant. Journal of Chemical Ecology 11: 1349–1358.

Bravo-Monzón AE, Ríos-Vásquez E, Delgado-Lamas G, Espinosa-García FJ. 2014. Chemical diversity among populations of *Mikania micrantha*: geographic mosaic structure and herbivory. Oecologia 174: 195–203.

Bustos-Segura C, Poelman EH, Reichelt M, Gershenzon J, Gols R. 2017. Intraspecific chemical diversity among neighbouring plants correlates positively with plant size and herbivore load but negatively with herbivore damage (C Scherber, Ed.). Ecology Letters 20: 87–97.

Cao Y, Jiang T, Girke T. 2008. A maximum common substructure-based algorithm for searching and predicting drug-like compounds. Bioinformatics 24: i366–i374.

Cao P, Yang J, Xia L, Zhang Z, Wu Z, Hao Y, Liu P, Wang C, Li C, Yang J, et al. 2024. Two gene clusters and their positive regulator SlMYB13 that have undergone domestication-associated negative selection control phenolamide accumulation and drought tolerance in tomato. Molecular Plant 17: 579–597.

Cereto-Massagué A, Ojeda MJ, Valls C, Mulero M, Garcia-Vallvé S, Pujadas G. 2015. Molecular fingerprint similarity search in virtual screening. Methods 71: 58–63.

Cosmo LG, Yamaguchi LF, Felix GMF, Kato MJ, Cogni R, Pareja M. 2021. From the leaf to the community: Distinct dimensions of phytochemical diversity shape insect–plant interactions within and among individual plants (T Züst, Ed.). Journal of Ecology 109: 2475–2487.

Diethelm AC, Reichelt M, Pringle EG. 2024. Herbivores disrupt clinal variation in plant responses to water limitation. Journal of Ecology 112: 338–347.

Dyer LA. 2018. Multidimensional diversity associated with plants: a view from a plant-insect interaction ecologist. American Journal of Botany 105: 1439–1442.

Ernst M, Kang KB, Caraballo-Rodríguez AM, Nothias L-F, Wandy J, Chen C, Wang M, Rogers S, Medema MH, Dorrestein PC, et al. 2019. MolNetEnhancer: Enhanced Molecular Networks by Integrating Metabolome Mining and Annotation Tools. Metabolites 9: 144.

Etienne Laliberté et. al. 2014. FD: Measuring Functional Diversity (FD) from Multiple Traits, and Other Tools for Functional Ecology. : 1.0–12.3.

Forrister DL, Endara M, Soule AJ, Younkin GC, Mills AG, Lokvam J, Dexter KG, Pennington RT, Kidner CA, Nicholls JA, et al. 2023. Diversity and divergence: evolution of secondary metabolism in the tropical tree genus *Inga*. New Phytologist 237: 631–642.

Glassmire AE, Philbin C, Richards LA, Jeffrey CS, Snook JS, Dyer LA. 2019. Proximity to canopy mediates changes in the defensive chemistry and herbivore loads of an understory tropical shrub, *Piper kelleyi* (T Turlings, Ed.). Ecology Letters 22: 332–341.

Herzler M, Herre S, Pragst F. 2003. Selectivity of substance identification by HPLC-DAD in toxicological analysis using a UV spectra library of 2682 compounds. Journal of Analytical Toxicology 27: 233–242.

Iason G, O’Reilly-Wapstra J, Brewer M, Summers R, Moore B. 2011. Do multiple herbivores maintain chemical diversity of Scots pine monoterpenes? Philosophical Transactions of the Royal Society B: Biological Sciences 366: 1337–1345.

Junker RR. 2018. A biosynthetically informed distance measure to compare secondary metabolite profiles. Chemoecology 28: 29–37.

Kessler A, Kalske A. 2018. Plant secondary metabolite diversity and species interactions. Annual Review of Ecology, Evolution, and Systematics 49: 115–138.

Kim HW, Wang M, Leber CA, Nothias L-F, Reher R, Kang KB, Van Der Hooft JJJ, Dorrestein PC, Gerwick WH, Cottrell GW. 2021. NPClassifier: A deep neural network-based structural classification tool for natural products. Journal of Natural Products 84: 2795–2807.

Li D. 2018. hillR: taxonomic, functional, and phylogenetic diversity and similarity through Hill numbers. Journal of Open Source Software 3: 1041.

Li D, Gaquerel E. 2021. Next-generation mass spectrometry metabolomics revives the functional analysis of plant metabolic diversity. Annual Review of Plant Biology 72: 867–891.

Liu X, Vrieling K, Klinkhamer PGL. 2017. Interactions between plant metabolites affect herbivores: A study with pyrrolizidine alkaloids and chlorogenic acid. Frontiers in Plant Science 8: 903.

Marion ZH, Fordyce JA, Fitzpatrick BM. 2015. Extending the concept of diversity partitioning to characterize phenotypic complexity. The American Naturalist 186: 348–361.

Mundim FM, Pringle EG. 2018. Whole-plant metabolic allocation under water stress. Frontiers in Plant Science 9: 852.

Nguyen D, D’Agostino N, Tytgat TOG, Sun P, Lortzing T, Visser EJW, Cristescu SM, Steppuhn A, Mariani C, Van Dam NM, et al. 2016. Drought and flooding have distinct effects on herbivore-induced responses and resistance in *Solanum dulcamara*. Plant, Cell & Environment 39: 1485–1499.

Nguyen D, Poeschl Y, Lortzing T, Hoogveld R, Gogol-Döring A, Cristescu SM, Steppuhn A, Mariani C, Rieu I, Van Dam NM. 2018. Interactive responses of *Solanum dulcamara* to drought and insect feeding are herbivore species-specific. International Journal of Molecular Sciences 19: 3845.

Onkokesung N, Gaquerel E, Kotkar H, Kaur H, Baldwin IT, Galis I. 2012. MYB8 controls inducible phenolamide levels by activating three novel hydroxycinnamoyl-coenzyme A: polyamine transferases in *Nicotiana attenuata*. Plant Physiology 158: 389–407.

Petrén H, Anaia RA, Aragam KS, Bräutigam A, Eckert S, Heinen R, Jakobs R, Ojeda-Prieto L, Popp M, Sasidharan R, et al. 2024. Understanding the chemodiversity of plants: Quantification, variation and ecological function. Ecological Monographs 94: e1635.

Petrén H, Köllner TG, Junker RR. 2023. Quantifying chemodiversity considering biochemical and structural properties of compounds with the R package CHEMODIV. New Phytologist: nph.18685.

Richards LA, Dyer LA, Forister ML, Smilanich AM, Dodson CD, Leonard MD, Jeffrey CS. 2015. Phytochemical diversity drives plant–insect community diversity. Proceedings of the National Academy of Sciences 112: 10973–10978.

Salazar D, Lokvam J, Mesones I, Vásquez Pilco M, Ayarza Zuñiga JM, de Valpine P, Fine PVA. 2018. Origin and maintenance of chemical diversity in a species-rich tropical tree lineage. Nature Ecology & Evolution 2: 983–990.

Sasidharan R, Grond SG, Champion S, Eilers EJ, Müller C. 2024. Intraspecific plant chemodiversity at the individual and plot levels influences flower visitor groups with consequences for germination success. Functional Ecology 38: 2665–2678.

Sedio BE. 2017. Recent breakthroughs in metabolomics promise to reveal the cryptic chemical traits that mediate plant community composition, character evolution and lineage diversification. New Phytologist 214: 952–958.

Steppuhn A, Baldwin IT. 2007. Resistance management in a native plant: nicotine prevents herbivores from compensating for plant protease inhibitors. Ecology Letters 10: 499–511.

Suzuki N, Rivero RM, Shulaev V, Blumwald E, Mittler R. 2014. Abiotic and biotic stress combinations. New Phytologist 203: 32–43.

Volf M, Segar ST, Miller SE, Isua B, Sisol M, Aubona G, Šimek P, Moos M, Laitila J, Kim J, et al. 2018. Community structure of insect herbivores is driven by conservatism, escalation and divergence of defensive traits in *Ficus* (T Turlings, Ed.). Ecology Letters 21: 83–92.

Wang Y, Backman TWH, Horan K, Girke T. 2013. fmcsR: mismatch tolerant maximum common substructure searching in R. Bioinformatics 29: 2792–2794.

Wang M, Carver JJ, Phelan VV, Sanchez LM, Garg N, Peng Y, Nguyen DD, Watrous J, Kapono CA, Luzzatto-Knaan T, et al. 2016. Sharing and community curation of mass spectrometry data with global natural products social molecular networking. Nature Biotechnology 34: 828–837.

Wetzel WC, Whitehead SR. 2020. The many dimensions of phytochemical diversity: linking theory to practice (H Hillebrand, Ed.). Ecology Letters 23: 16–32.

Whitehead SR, Bass E, Corrigan A, Kessler A, Poveda K. 2021. Interaction diversity explains the maintenance of phytochemical diversity (R Bardgett, Ed.). Ecology Letters 24: 1205–1214.

Ziaja D, Müller C. 2023. Intraspecific chemodiversity provides plant individual- and neighbourhood- mediated associational resistance towards aphids. Frontiers in Plant Science 14: 1145918.

